# Mechanical, physical and chemical characterisation of mycelium-based composites with different types of lignocellulosic substrates

**DOI:** 10.1101/569749

**Authors:** Elise Elsacker, Simon Vandelook, Joost Brancart, Eveline Peeters, Lars De Laet

## Abstract

The current physical goods economy produces materials by extracting finite valuable resources without taking their end of the life and environmental impact into account. Modernity leaves us with devasted landscapes of depleted resources, waste landfill, queries, oil platforms. At the time of the Anthropocene, the various effects the human role has on the constitution of the soils create an acceleration of material entropy. It is the terrestrial entanglement of fungal materials that we investigate in this paper by offering an alternative fabrication paradigm based on the growth of resources rather than on extraction. Unlike the latter, biologically augmented building materials can be grown by combining micro-organisms such as fungal mycelium with agricultural plant-based waste. In this study, we investigate the production process, the mechanical, hygrothermal and chemical properties of mycelium-based composites with different types of lignocellulosic reinforcement fibres combined with a white rot fungus, *Trametes versicolor*. Together, they form an interwoven three-dimensional filamentous network binding the feedstock into a lightweight material. The mycelium-based material is heat-killed after the growing process. This is the first study reporting the dry density, the Young’s modulus, the compressive stiffness, the stress-strain curves, the thermal conductivity, the water absorption rate and a complete FTIR analyse of mycelium-based composites by making use of a disclosed protocol with *T. versicolor* and five different type of fibres (hemp, flax, flax waste, softwood, straw) and fibre conditions (loose, chopped, dust, pre-compressed and tow). These experimental results show that mycelium-composites can fulfil the requirements of thermal insulation. The thermal conductivity and water absorption coefficient of the mycelium composites with flax, hemp, and straw have an overall good insulation behaviour in all the aspects compared to conventional unsustainable materials. The conducted tests reveal that the mechanical performances of the mycelium-based composites depend more on the fibre condition, size, and processing than on the chemical composition of the fibres.

Graphical abstract

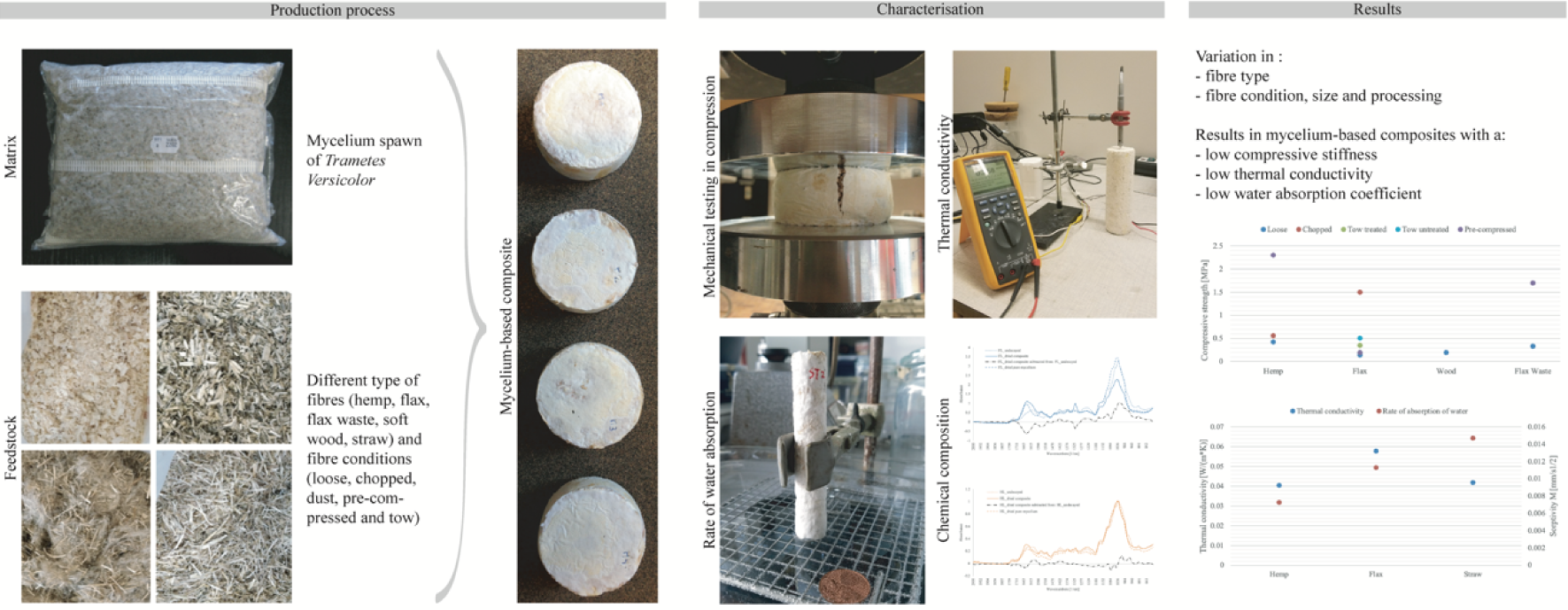

**Highlights:**

- The type of fibre influences the colonisation of mycelium: samples containing flax, hemp, straw and flax-waste resulted in a well-developed composite
- The type of fibre has a smaller influence on the compressive stiffness than the fibre processing and size.
- Pre-compression and chopped fibres (particle size <5mm) improve the compressive mechanical properties of mycelium composites.
- The thermal conductivity and water absorption coefficient of the mycelium composites with flax, hemp, and straw have an overall good insulation behaviour in all the aspects compared to conventional unsustainable materials.

## 1. Introduction

The physical goods economy requires the mining of raw resources and turns large quantities of those goods into trash at the end of the use. A linear material economy is destructive to the environment and causes pollution [1–4]. Already since the mid-sixties, scientist Kenneth E. Boulding pointed out the urge of facing the problems related to an increase in material entropy created by those activities [5]. Yet, on the first of August 2018, Earth’s annual resources budget has been consumed in just 7 months by human activity [6]. Our industrial society consume more natural resources than nature can regenerate by overfishing, overharvesting of forests, and emitting more carbon dioxide into the atmosphere than ecosystems can absorb. The construction sector in Europe accounts for about half of all our extracted materials and energy consumption and about a third of our water consumption and waste generation [7].

To meet the limits of our planet, biologically augmented building materials can be grown with agricultural plant-based waste instead of extracting raw resources that will generate future waste. The closed-loop materials can generate new materials, new life from decay. The renewable mycelium-based composites are composed of fungal biomass, called mycelium and lignocellulosic waste streams. The hyphae of the fungus form an interwoven three-dimensional filamentous network through the cellulose, hemicellulose and lignin rich substrate by digesting its nutrients and simultaneously binding the substrate. When reaching complete substrate colonization the organism is heat-killed above a critical temperature to render the material inert and allow the evaporation of all residual water from the material. The result is a lightweight and bio-degradable composite with a low impact on the environment. This material has the potential to replace fossil-based and synthetic materials such as polyurethane and polystyrene.

The potential benefits of this material in the field of architecture and construction have hardly been investigated so far [8]. Additionally, the composite material is complex due to its wide variety of possible combinations between substrate type and fungal species. Studies on the effects between the process variables and the material behaviour are very limited. Every growth parameter variation can result in changes in the material constitution and mechanical properties. The main factors affecting the production of mycelium composites, and consequently their mechanical behaviour, are: the matrix (mycelium species), the feedstock selection (lignocellulosic substrate), the interaction between white rot fungi and their feedstock and last but not least the process variables during manufacturing (protocol, sterilisation, inoculation, packing, incubation, growing period and drying method). Studies have demonstrated that the mechanical properties of mycelium-composites are affected by the feedstock [9–11]. The average compressive strength of cotton down woven mat and hemp pitch woven and non-woven mat substrates varies between 0,67 to 1.18 MPa at 60% deformation in sample height [11]. Substrates containing not degraded natural fibres exhibit a strain-hardening behaviour because the natural fibres serve to reinforce and prevent shear failure [12]. The average flexural modulus is respectively 1 MPa, 3 MPa and 9 MPa for non-pressed cotton, rapeseed and beechwood based composites. For heat pressed composites the flexural modulus is respectively 34 MPa, 72 MPa and 80 MPa [9]. The average flexural modulus ranged from 66.14 to 71.77 MPa for cotton down woven mat, hemp pith woven and non-woven mat [11]. The packing method influences the thermal conductivity of mycelium-based composites. Loosely packed substrate has a low thermal conductivity of 0,05 W/(m · K), whereas densely packed substrate has a slightly higher thermal conductivity of 0,07 W/(m · K), due to less dry-air cavities [12]. The water absorption properties are reported to be high, between 298% and 350% [11]. However, the methodologies, mycelium species and feedstocks of the published studies proceed in no standardised and comparable way. Moreover, many studies do not fully disclose the preparation of the mycelium-composites due to proprietary information preventing a proper comparison or replication [9,13–15].

The aim of this study is to present a comprehensive overview of the production processes, the mechanical, hygrothermal and chemical properties of mycelium-based composites grown on different lignocellulosic substrate types. The fibres that were considered for the characterisation included flax, flax waste, flax dust, untreated flax tow, treated flaw tow, wood, hemp, straw and straw dust. *Trametes (Coriolus) versicolor*, (L.) Lloyd (1921), was the selected fungus species for all substrate types. It is also commonly known as Turkey tail, a widespread polypore basidiomycete belonging to the white-rot fungi that inhabits dead tree trunks.

The results presented in this paper cover the dry density, the Young’s modulus, the compressive stiffness, the stress-strain curves, the thermal conductivity, and the water absorption rate of the mycelium-based composites. In addition, a Fourier-transform infrared spectroscopy (FTIR) analyse was conducted on undecayed fibres, dried decayed fibres, and pure mycelium to relate the mechanical results of mycelium-composites to the breakdown of lignin, hemicellulose and cellulose by *Trametes versicolor*.

## 2. Materials and methods

### 2.1. Materials

The variations in the natural fibres are based on two different aspects, the type of fibre and the fibre condition and size (loose, chopped, dust, pre-compressed and tow). The used fibres are summarized in Table 1. The flax, the flax dust, the flax long treated fibres, the flax long untreated fibres, the flax waste, the hay dust and the straw were obtained from Jopack bvba (Rumbeke, Belgium). The hemp fibres and pine softwood fibres were purchased from Aniserco S.A (Groot-Bijgaarden, Belgium). The particle size of loos fibres varied between 10 and 5mm. Tow fibres were cut to a length of 100mm. The treated fibres are very similar in appearance to the untreated fibres but are slightly softer to the touch and ready for spinning to yarn, while the untreated fibres had more residual small fibres. Chopped fibres where processed in a blender to reach a fibres size smaller than 5 mm. Dust fibres had a particle size smaller than 0,5 mm.

**Table 1:**
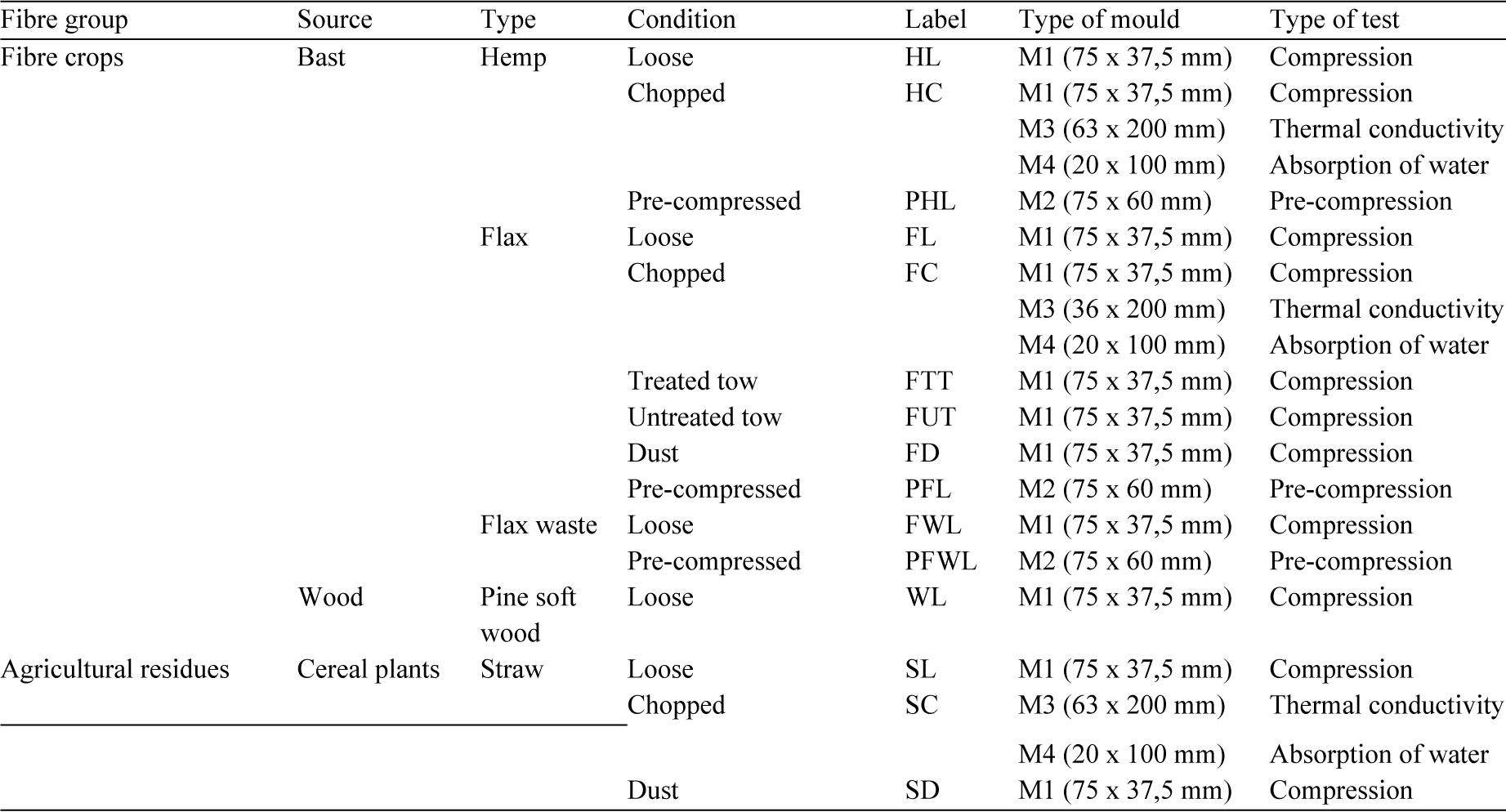
Summary of the natural fibre types, conditions, type of mould and type of test, with the corresponding labels.

The mycelium spawn *Trametes versicolor* (M9912-5LSR-2 O447A) was purchased from Mycelia bvba (Nevele, Belgium). It was conserved on a grain mixture at 4°C in a breathing Microsac bag (Sac O2 nv, Nevele, Belgium) of 5 litres.

### 2.2. Sample preparation

#### 2.2.1. Fibre preparation

The used fibres vary in size. To have the adequate fibre size for the tests, the fibres were soaked in water during 24 hours. They then were rinsed abundantly, after which they were placed in a blender during 10 minutes with fresh water. The fibres were sieved with a 5mm strainer, squeezed manually and spread on a plate in a warm room to allow the water to evaporate. The dried chopped fibres were then transferred in an autoclavable micro box (obtained from Sac O2 nv). The loose, tow and dust fibres were not processed and directly transferred to a micro box. A small cotton piece was placed between the cover and the box to achieve the same pressure inside and outside the box while autoclaving. Additionally, a silver foil was used to cover the box in order to preserve the filter on the cover and avoid water infiltration inside the box during autoclaving. The fibres were sterilised to render the substrate inert. This was done by placing the closed boxes in an autoclave machine during 20 minutes at 121°C. The boxes were left to cool down for 24 hours.

#### 2.2.2. Moulds

The moulds for the compression test were fabricated from a hollow PVC tube that is composed of two demountable parts. The geometry of the moulds depends from the type of test. As no standard testing procedure exists, different geometries were tested in a first iteration based on literature [12,14,16,17]. Eventually a diameter to height ratio of 2:1 was selected for compressive tests; the test samples have a diameter of 75 mm and a height of 37,5 mm (M1). The large contact area with the loading bench resulted in a more distributed stress induction, while the limited height prevented failure by buckling. For pre-compressed samples, moulds of diameter 75 mm and height 60 mm (M2) were used.

The PVC moulds that were fabricated for the thermal conductivity tests, were based on [18], [19] and [20]. The moulds had a diameter of 63 mm and height of 200 mm (M3). The same samples were used to determine the elastic and damping properties of the material according to [21].

To define the rate of absorption of water by partial immersion the norms [22] and [23] were applied, and PVC moulds of 20 mm in diameter and 100 mm height were used (M4).

### 2.3. Composite fabrication

#### 2.3.1. Inoculation

The inoculation is performed in sterile conditions inside a laminar flow. The fibres, sterile H_2_O and spawn necessary to fill the moulds are weighted, mixed together and put in the moulds. The added water is three times the weight of the fibres. Then, 10% spawn is added to the substrate. The moulds are filled by layers, while compressing each layer with a spoon to obtain a compact and dense sample (Figure 2a). A transparent foil or parafilm is used to seal the moulds.

**Figure 1:**
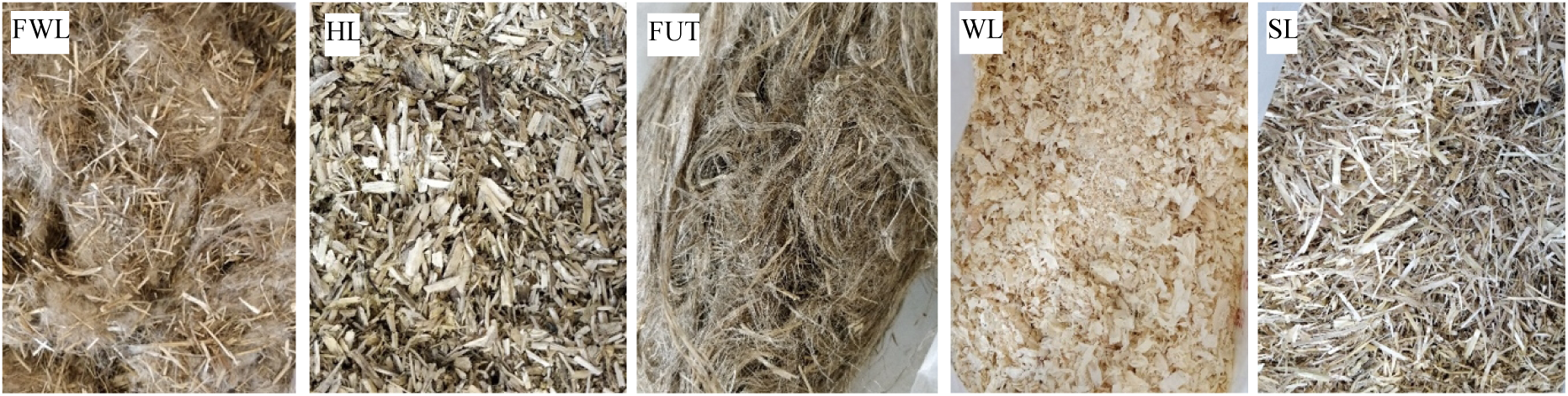
Fibre types loose flax waste (FWL), loose hemp (HL), flax untreated tow (FUT), loose wood (WL), loose straw (SL).

**Figure 2:**
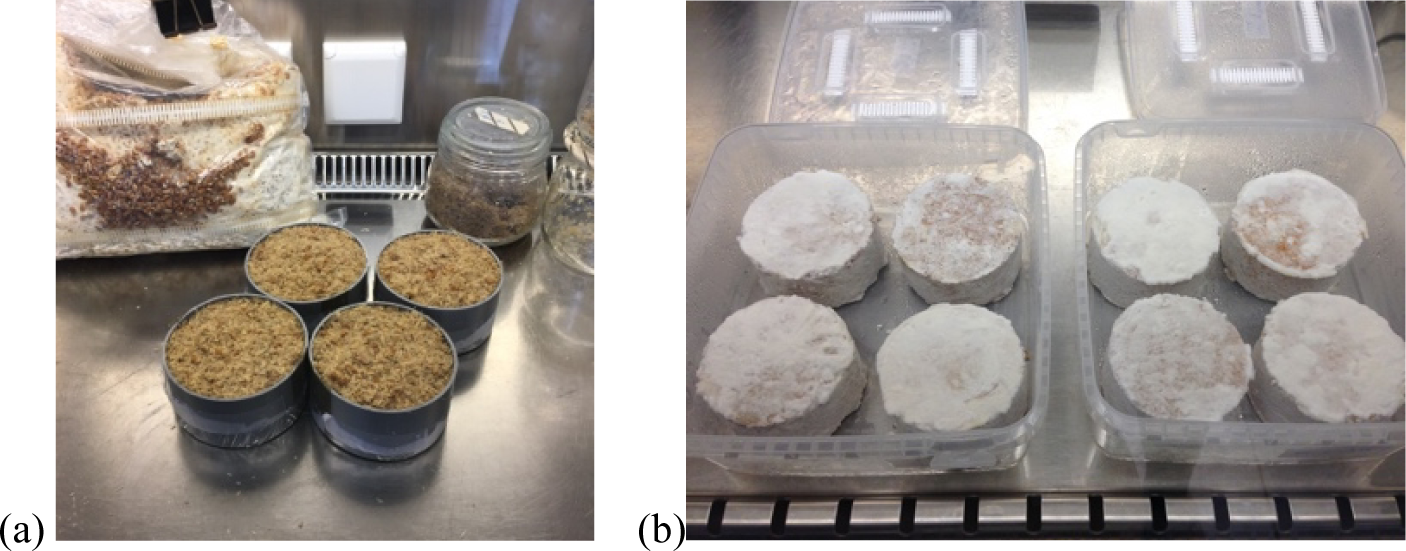
(a) Preparation of samples in moulds for the compression test in the laminar flow. (b) Unmoulding of samples after 8 days.

#### 2.3.2. Incubation

Samples were placed in a micro box to create a sterile micro climate. Finally, the filled boxes were placed in an incubator at 28°C. Since the PVC mould only allows gas exchange at bottom and top of the moulds, a second growth phase was introduced to enable mycelium to grow throughout the entire sample. Therefore, after 8 days, the samples were unmoulded in the laminar flow (Figure 2b) and incubated in a microbox for another minimum of 8 days without mould. The growth periods of every type of sample are different and indicated in Table 2.

**Table 2:**
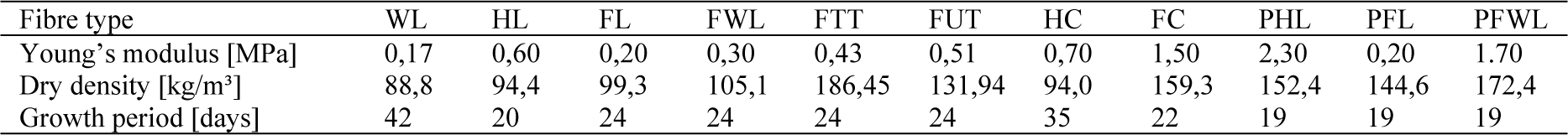
Summary of the compressive Young’s modulus, dry density and growth period

| Fibre type | WL | HL | FL | FWL | FTT | FUT | HC | FC | PHL | PFL | PFWL |
| --- | --- | --- | --- | --- | --- | --- | --- | --- | --- | --- | --- |
| Young's modulus [MPa] | 0,17 | 0,60 | 0,20 | 0,30 | 0,43 | 0,51 | 0,70 | 1,50 | 2,30 | 0,20 | 1,70 |
| Dry density [kg/m<sup>3</sup>] | 88,8 | 94,4 | 99,3 | 105,1 | 186,45 | 131,94 | 94,0 | 159,3 | 152,4 | 144,6 | 172,4 |
| Growth period [days] | 42 | 20 | 24 | 24 | 24 | 24 | 35 | 22 | 19 | 19 | 19 |

The PHL, PFL, PFWL samples were pre-compressed after 8 days. Samples were placed on a base plate that was fixed on a table. Every sample was provided with a cover as surface barrier between the sample and the screw clamp. Two screw clamps were required for every sample, one to compress and one to keep the mould closed (Figure 3). Samples remained in this position during the second growth phase.

**Figure 3:**
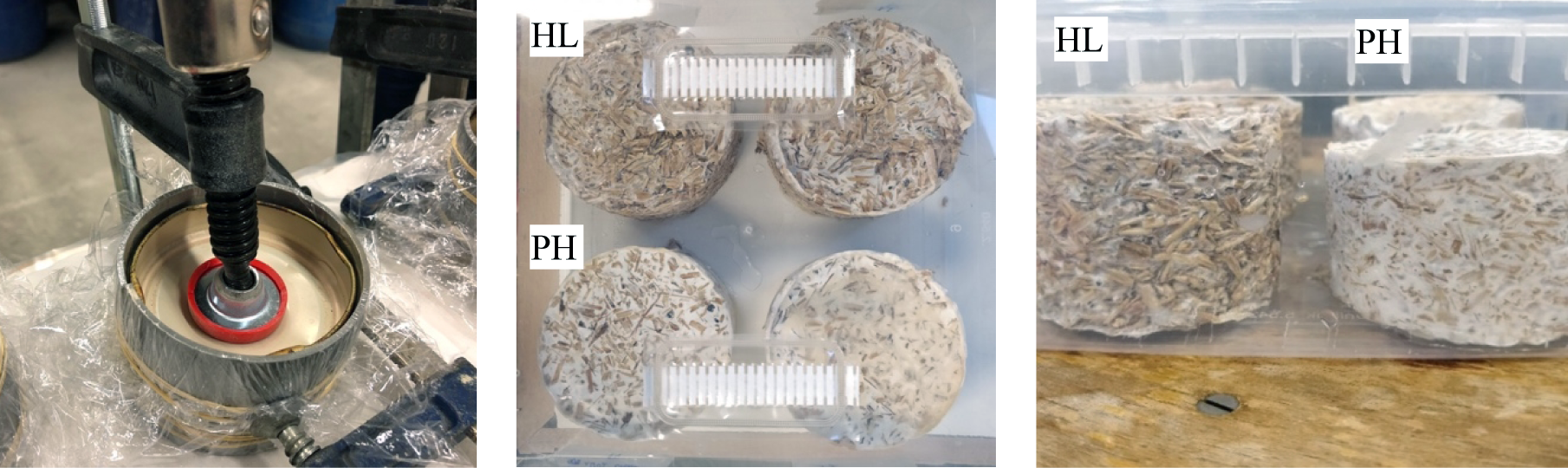
Pre-compressed PHL, PFL, PFWL samples. Unmoulded HL (above, right) and PHL (under, left) samples after 8 days at 28°C, the second growth phase is initiated in a sterile microbox with filters.

**Figure 4:**
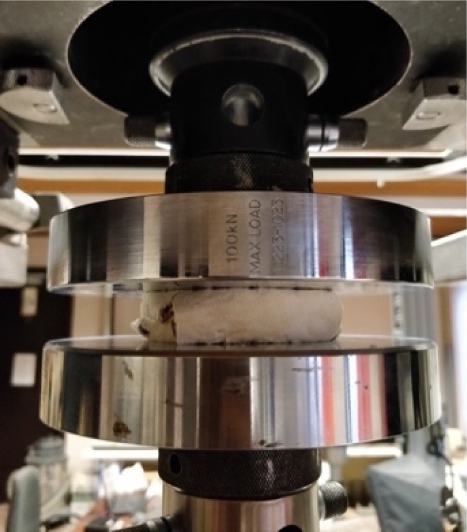
Instron load bench for compression tests at the state of maximum deformation of the FWL sample.

#### 2.3.3. Drying process

To render the material inert, all samples were dried in a convection oven at a temperature of 70°C for 5 to 10 hours, until their weight stabilised and thus all water was evaporated. The dry weight, dry density and the moisture content, as well as all dimensions and shrinkage were calculated for every sample and are available upon request to the corresponding author.

### 2.4. Mechanical testing in compression

The compressive stiffness was obtained, based on ASTM D3501 [24], on an Instron 5900R load bench, at ambient conditions (25°C and ∼ 50%), with a capacity of 100 kN and a load cell of 10 kN. The 10 kN load cell is used to have the most accurate results since low values of ultimate loads are to be expected. The tests were displacement controlled with a rate of 5 mm/min. The contact surface was not perfect due to the rough surfaces of the samples. The test was stopped when a fixed strain was reached in the specimen, varying between 70% and 80%. The load-displacement curve was converted to a stress-strain curve, using the following formulas to calculate the compressive stress σ and the strain ε:

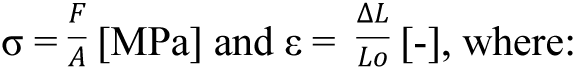

F = compressive force [N], A = original cross section of the specimen [mm²], ΔL = displacement of the loading surfaces [mm] and L_o_ = original height of the test piece [mm].

### 2.5. Thermal conductivity

The transient method was used to measure the thermal conductivity according to ASTM D 5334 – 00. The thermal needle probe (produced by Huksefluxand with commercial reference TP02), complies with the standard. Other equipment needed were: a constant current power source, thermal reading tool (multimeter), voltage reader, stopwatch or software to program the time steps of reading (fluke software), driller or stiff needle to open the guide hole in the middle for the probe. A first guide hole was drilled in the centre of the cylindric sample with the help of an extra needle with a smaller diameter than the TNP. There is one exception from the ASTM standard: a different electrical current (Table 3) in the circuit was applied.

**Table 3:** Summary of thermal conductivity, density and moisture content of chopped flax (FC), hemp (HC) and straw (SC) mycelium composites.

| Fibre type | Current in TNP [A] | Density [Kg/m<sup>3</sup>] | Thermal conductivity ($\lambda$) [W/ (m*K)] | Average ($\lambda$) [W/ (m*K)] | Moisture content (%) |
| --- | --- | --- | --- | --- | --- |
| FC_B1 | 0.104 | 134.71 | 0.0582 | 0.0578 | 9,5466 |
| FC_B2 | 0.071 |  | 0.0603 |  |  |
| FC_B3 | 0.175 |  | 0.0550 |  |  |
| HC_B1 | 0.106 | 98.92 | 0.0409 | 0.0404 | 7,3795 |
| HC_B2 | 0.177 |  | 0.0417 |  |  |
| HC_B3 | 0.088 |  | 0.0386 |  |  |
| SC_B1 | 0.103 | 94.39 | 0.0420 | 0.0419 | 12,9625 |
| SC_B2 | 0.135 |  | 0.0421 |  |  |
| SC_B3 | 0.173 |  | 0.0417 |  |  |

### 2.6. Rate of water absorption

Among the different ways to determine the capillarity of construction materials, the method for Measurement of Rate of Absorption of Water by Hydraulic-Cement Concretes ASTM C 1585 with partially submerged samples, was applied to the mycelium composites because this method is commonly used, easily applicable, has reliable results, and it is a non-destructive test. The tests were conducted in stable lab conditions with environmental temperature changes lower than 3°C, and the humidity changes in the air lower than 5%.

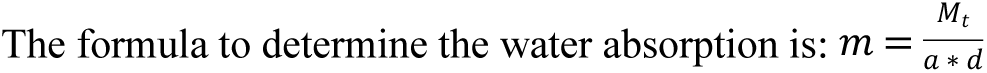

With m: water absorption [mm], M_t_ = is the change of mass of the specimen [g], a = the exposed cross-section area [mm^2^], d= the density of the water [g/mm^3^]

### 2.7. Chemical characterization

For the Fourier Transform Infrared Spectroscopy all IR spectra are acquired on a Nicolet 6700 FT-IR spectrometer from Thermo Fischer Scientific. The FTIR instrument is equipped with an IR source, DGTS KBr detector and KBr beamsplitters and windows. FTIR spectra are recorded in single bounce Attenuated Total Refractance (ATR) mode using the Smart iTR accessory, equipped with a diamond plate (42° angle of incidence). The spectra are recorded with automatic atmospheric correction for the background.

All samples were measured at a spectral resolution of 4 cm^−1^ with 64 scans per sample. Peak height, area and subtraction was measured using Spectragryph v1.2.8 software. The spectra were cut between 4000 and 700 cm^−1^ bands, then the baseline was constructed by connecting the lowest data point on either side of the peak, and finally the peaks were normalised by surface area. The maximum absorbance intensities for lignin associated bands were divided against carbohydrate reference peaks in Excel.

## 3. Results and discussion

### 3.1. Sample description and growth examination

In parallel to the composite fabrication in moulds, samples were grown in squared petri-dishes (120 x 120 x 17mm) to follow the growth day by day. The growth evolution of a representative selection of the samples is presented in Figure 5. The samples with flax dust (FD), pine softwood (W) and straw (S) were poorly grown after 10 days. Due to a slow growth all flax dust (FD) samples systematically contaminated. Therefore, no further test were conducted with this type of fibre. Samples containing pine softwood (W) and straw (S) ultimately grow. We can observe a dense white chitinous layer formed all over the hemp (HL), flaw (FL), flax waste (FWL) and flaw tow (FTT and FUT) specimen.

**Figure 5:**
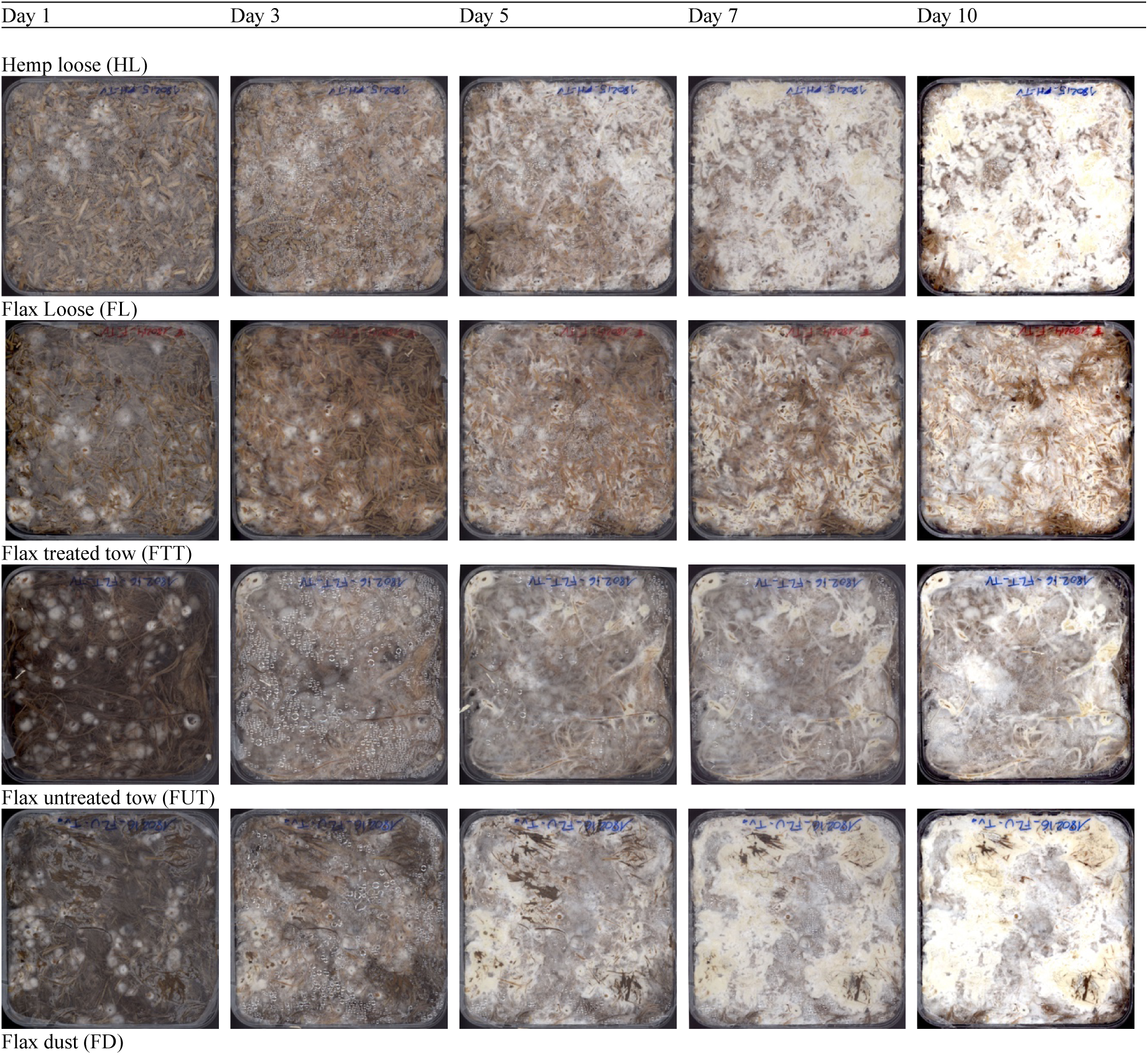
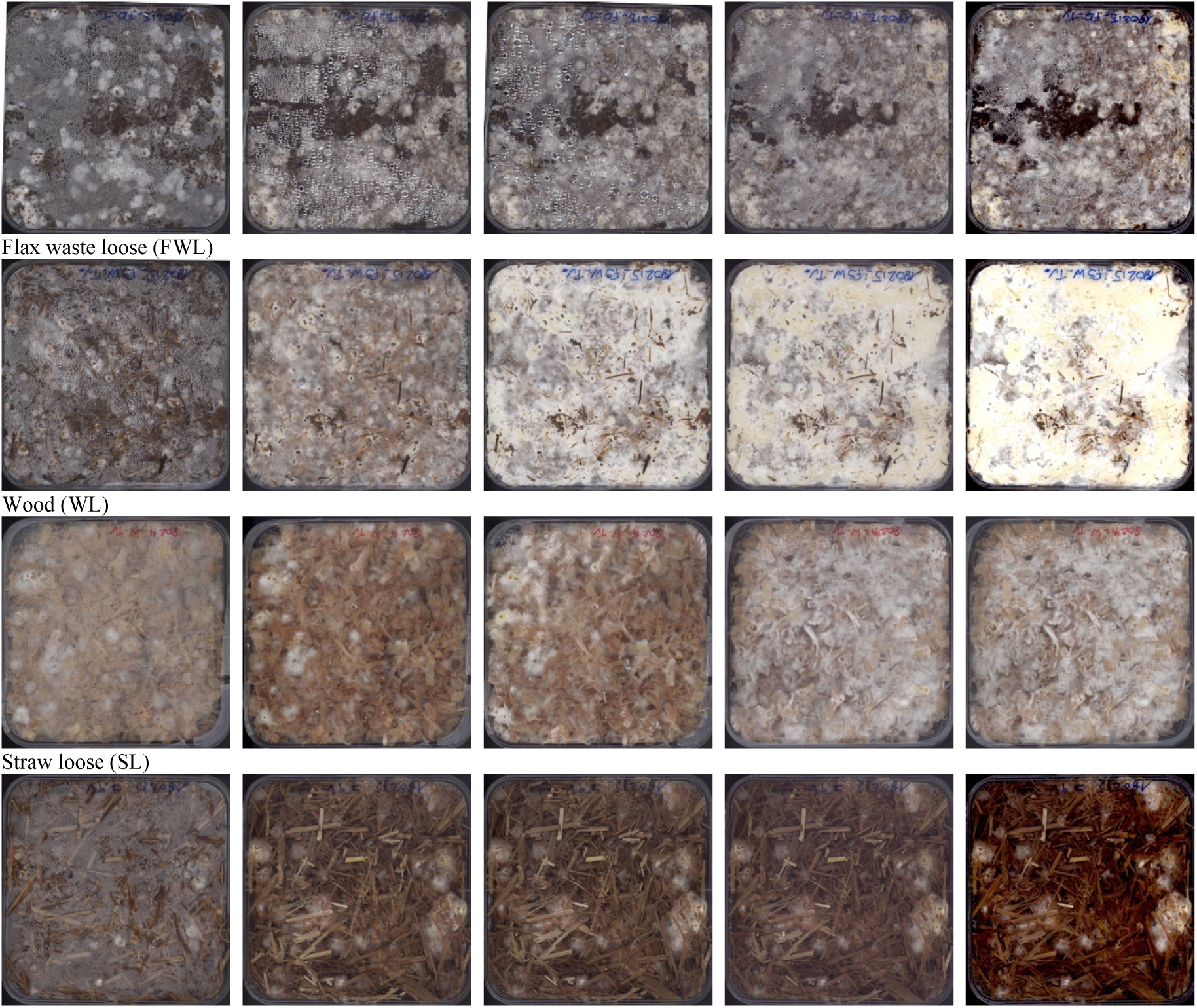
Growth evolution of a representative selection of the samples HL, FL, FTT, FUT, FD, FWL, WL, SL.

**Figure 6:**
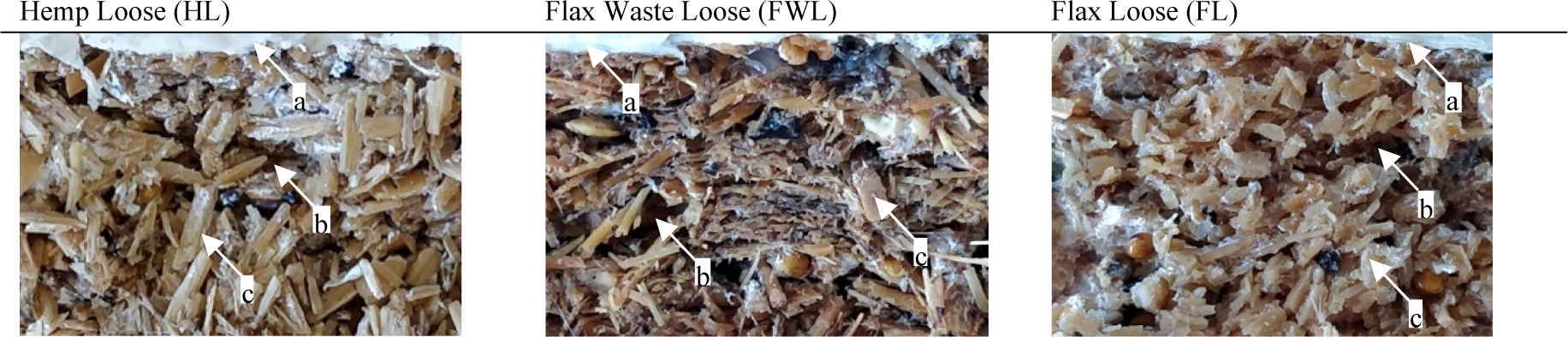
Cross section of the inner growth of undried FL, FWL and FL samples. (a) mycelium chitinous layer, (b) air-void, (c) limited decayed fibre by mycelium.

Undried samples of H, F and FW were cut open to analyse the inner growth of the specimens. Despite the spawn that was mixed throughout the whole substrate during inoculation, poor growth was observed inside the samples. A possible explanation for the low colonisation inside the sample might be the accumulation of heat produced by mycelium during growth, and the migration of hyphae to the surface of the samples, attracted by oxygen, after demoulding in the second growth phase. The outer surface grows, whereas internal growth does not continue.

The average moisture content of all living decayed samples before drying was 76,8 wt% (weight percentage). By controlling the weight of the samples during drying, their humidity was determined. The weight of the loose hemp samples decreased faster compared to the loose flax samples. The highest decrease in weight took place during the two first hours of drying, followed by a more constant behaviour for the third hour, after which weight decreased again until stability was reached after 5h (Figure 7).

**Figure 7:**
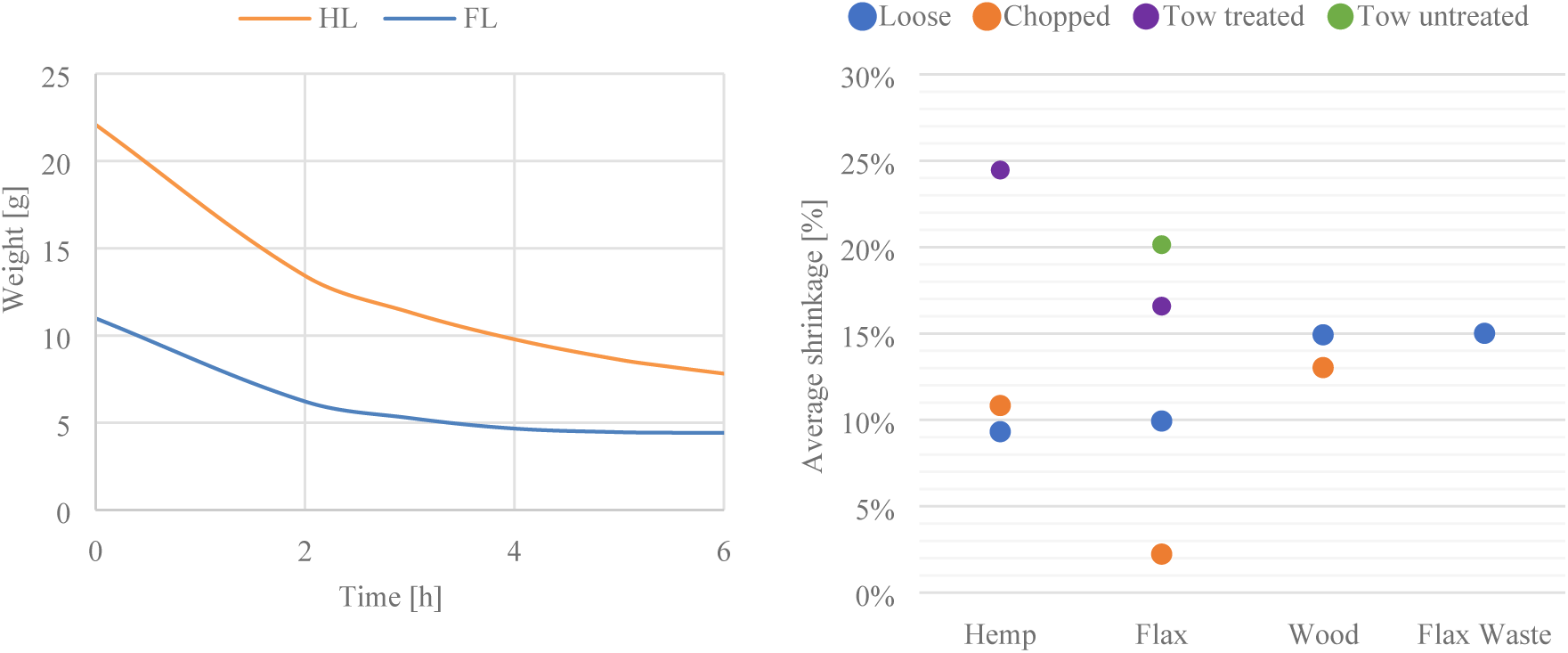
Left: drying curve for the HL and FL samples at a temperature of 80°C. Right: Average shrinkage percentage per fibre type and condition.

The samples were shrunk due to drying. For samples composed of loose hemp the average shrinkage percentage for diameter and height were respectively 7% and 12%. For loose flax, shrinkage percentages for diameter and height were respectively 6% and 14 %, and loose wood, respectively 10% and 20%. There was no noticeable difference in shrinkage between the fibre conditions: loose, chopped, tow. The average shrinkage percentage of all grown samples is 12% for the diameter and 14% for the height.

The dry density, summarize in Table 2 for all samples, was the lowest for loose flax with 59,77 kg/m^3^ and the highest for treated tow flax with 186,45 kg/m^3^.

### 3.2. Mechanical behaviour in compression

Within one group, all the loose fibre samples (WL, HL, FL, and FWL) behaved in a similar way, resulting in closely related stress-strain curves (Figure 8) for the compression test. The mechanical compressive stiffness was obtained from the slope of stress-strain curve with the tangent modulus and is summarized in Table 2 along with the average dry density and growth period. The values are in correspondence with the preceding growth observations; within the group of loose fibres the combination of mycelium and HL achieved the highest compressive stiffness, followed by FWL. The lowest stiffness is achieved by the WL samples, which also have the lowest density. We can also observe an increase in compressive stiffness for HC (0,70 MPa) and FC (1,50 MPa) compared to HL (0,60 MPa) and FL (0,20 MPa), indicating that the fibre condition and smaller fibre size influences the compressive stiffness. The compressive stiffness is considerably larger for FC, compared to any other fibre type and condition. This feature contradicts the previously discussed results for samples with a FL and HL substrate, where the Young’s modulus for samples with FL is three times smaller than the obtained value for HL. Chopped fibre substrates result in mycelium composites with slightly higher densities compared to the samples with a loose substrate. For HL and HC, the density remains approximately equal, which might explain the small change in stiffness. FC fibres increase the composite’s density, as well as its stiffness. Flax fibres have the highest stiffness in a chopped condition and the lowest one in a loose condition, whereas the results for hemp-based substrates are less spread. The compressive strength for FTT and FUT resulted in 0,43 MPa and 0,51MPa, which can be related to the higher density of the composite.

**Figure 8:**
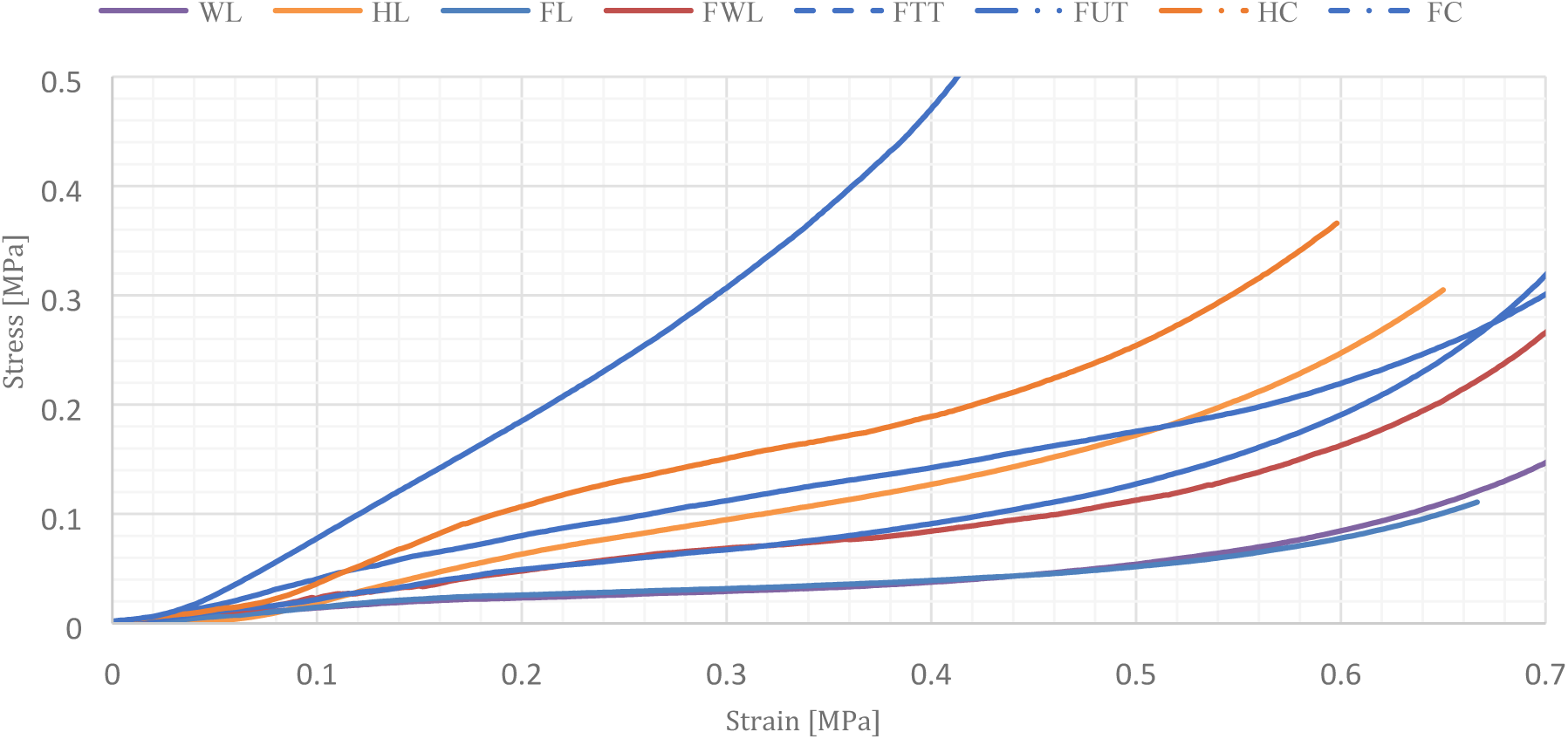
Stress-strain evolution of mycelium-composited with different types of fibres (WL, HL, FL, FWL, FTT, FUT, HC and FC) during uniaxial compression.

To understand the influence of the growth period on the compressive stiffness, an extra experiment was conducted with chopped flax. The FC samples with the shortest growth period (17 days) had a lower stiffness of 1.25 MPa, whereas the FC samples grown during 22 days had a stiffness of 1.50 MPa. In this case the growth period might have influenced the mechanical response of the composites. Those results should be confirmed by further research.

Additional pre-compressed samples PHL, PFL and PFWL were grown to optimize the compressive stiffness. Pre-compression of the samples took place during manufacturing, before demoulding and before drying. The aims was to improve the compactness and thus compressive properties of the composites. In addition to the compression of the material with a spoon during manufacturing, the material is compressed with screw clamps before initiating the second growing process. The results, shown in Figure 9 are obvious; the action of pre-compressing samples influences their mechanical response in terms of stiffness.

**Figure 9:**
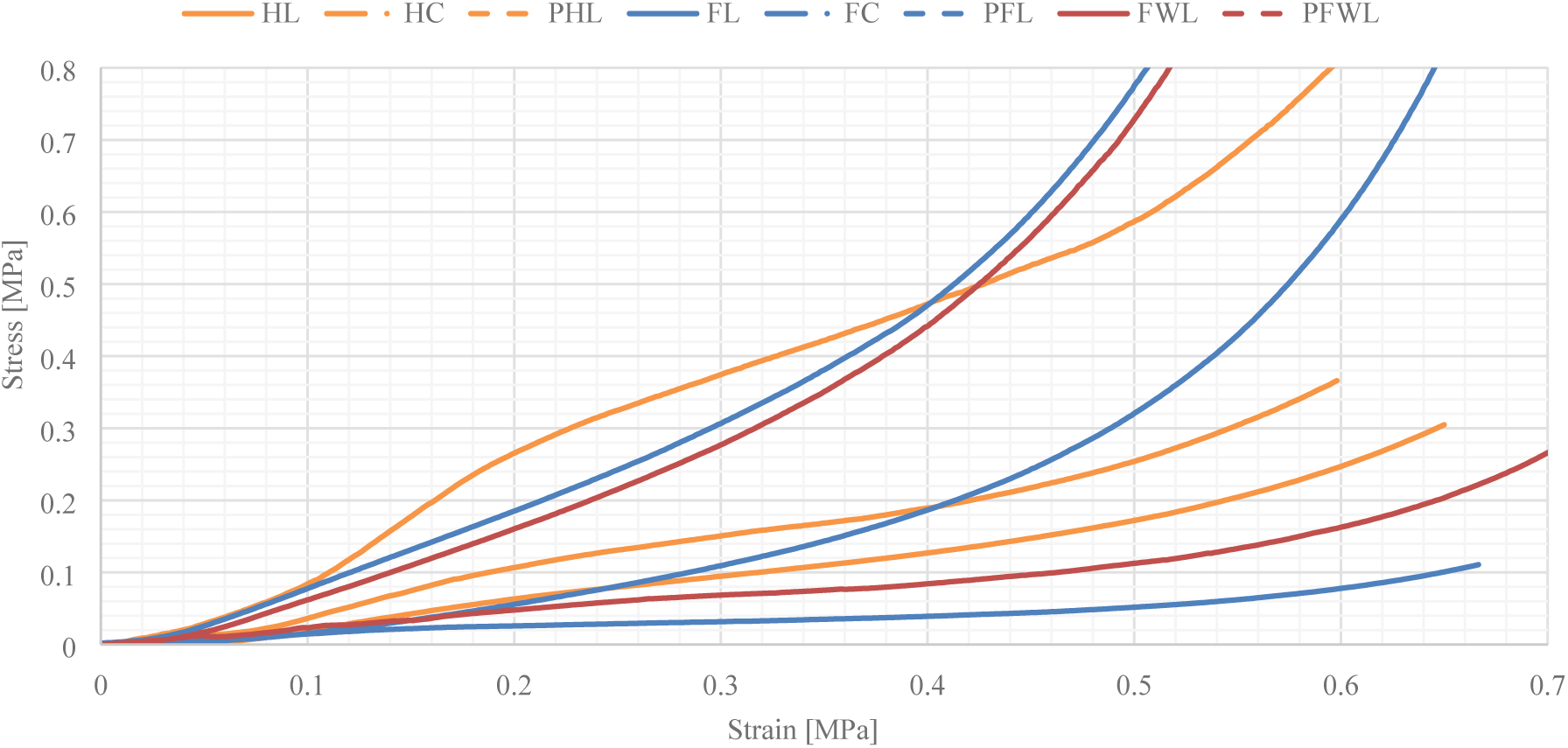
Effect of fibre condition (loose, chopped and pre-compressed) on the stress-strain response in compression. Solid lines indicate loose fibres, dot-dash lines indicate chopped fibres, and dashed lines indicate pre-compressed loose fibres.

### 3.3. Thermal conductivity

Table 3 presents the thermal conductivity of FC, HC and SC samples. The mycelium composites had a lower thermal conductivity than water, therefore lower electrical current was used to mark a clear increase of temperature during the test.

The temperature changes depending on the applied current during the test (Figure 10). The response of the probe’s temperature is monitored in function of the time [25]. Nonetheless, as expected, the overall results of the calculations did not present large differences. The higher the applied current, the faster the temperature increased, and thus the lower the thermal conductivity. Yet, this fast rise in temperature is not recommended by the standard [18] due to possible errors while readings. Therefore it was more reliable to take the average of the values for thermal conductivity corresponding to a lower applied current.

**Figure 10:**
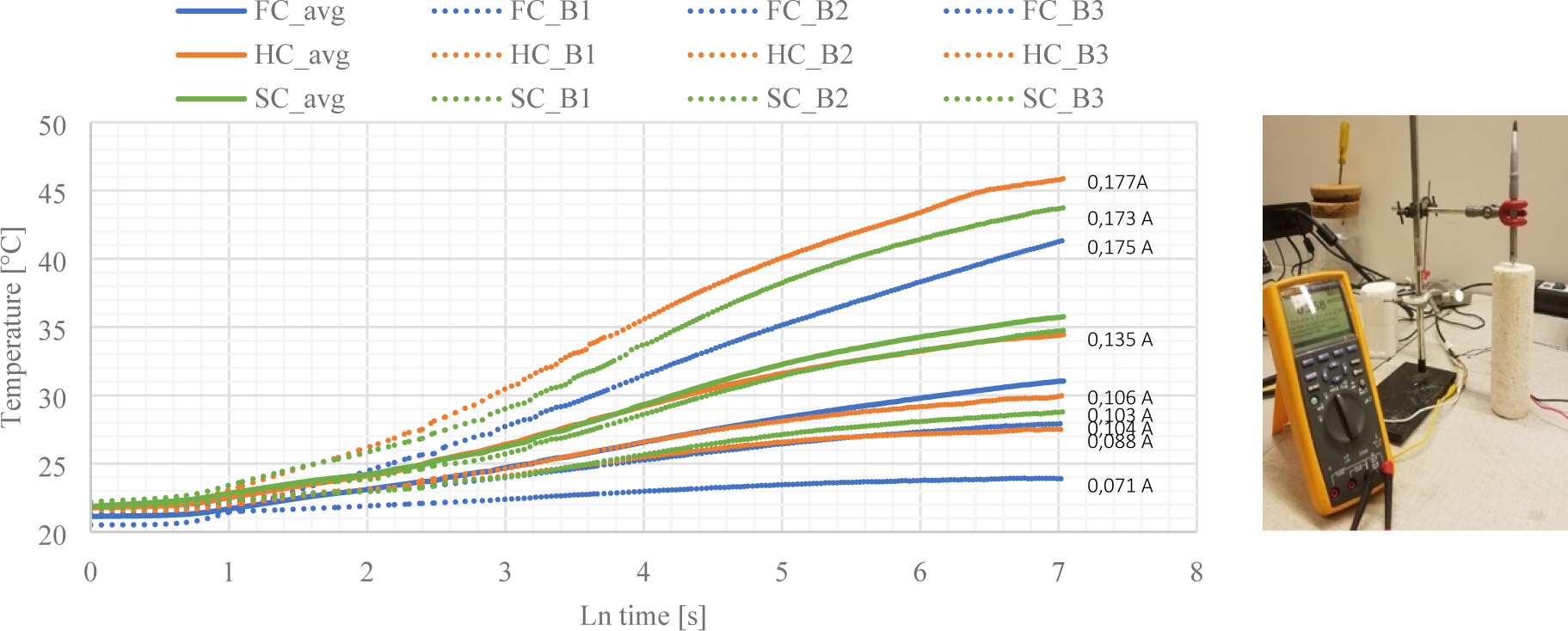
Temperature - time response of the thermal probes for the determination of the thermal conductivity of FC, HC and SC with different currents (A).

The value of the thermal conductivity for FC is 0.0578 W/ (m*K), which is larger than HC and SC, respectively 0.0404 W/ (m*K) and 0.0419 W/ (m*K). The larger thermal conductivity for FC is due to the higher density of the samples (134,71 kg/m^3^). The values of the thermal conductivity for HC and SC range within the same values as conventional insulating materials (Table 4) such as rock wool, glass fibres, sheep wool, cork. The results also show better thermal insulation properties compared to recent research on mycelium-based biofoams [12] with values between 0.05-0.07 W/ (m*K), and [26] with values between 0.078 and 0.081 W/ (m*K). This can be attributed to a different production protocol of the samples, species, different densities (180-380 kg/m^3^, and 51-62 kg/m^3^), and a different fibre type (sawdust pulp of Alaska birch, and wheat straw). The results prove that mycelium composites can become an alternative biological insulation material. However, other properties related to (the thermal performances of) insulating materials, such as fire resistance, aging, acoustics, water vapor diffusion, should be tested.

**Table 4:** Summary of thermal conductivity, density of mycelium-based composites and conventional insulation materials.

| Insulating Material | Density [Kg/m<sup>3</sup>] | Thermal conductivity [W/(m*K)] | Source |
| --- | --- | --- | --- |
| Mycelium-based composite | 94 - 135 | 0.0578 - 0.0404 |  |
| Rock wool | 470-2250 | 0.044 | [27] |
| Glass wool | 13-100 | 0.033-0.045 | [28] |
| Extruded polystyrene | 18-50 | 0.025-0.035 | [28] |
| Kenaf | 30-180 | 0.034-0.043 | [29] |
| Sheep wool plates | 10-25 | 0.038-0.054 | [29] |

**Table 5:** Summary of the water absorption rate of chopped flax (FC), hemp (HC), straw (SC).

| Fibre type | Water absorption rate M<br>[mm/s<sup>1/2</sup>] | Average water absorption rate M<br>[mm/s<sup>1/2</sup>] | Water absorbed after 24 hours<br>[%] | Density [Kg/m<sup>3</sup>] |
| --- | --- | --- | --- | --- |
| FC_B1 | 0.013 | 0.0113 | 30.28 | 136,43 |
| FC_B2 | 0.012 |  |  | 135,09 |
| FC_B3 | 0.0094 |  |  | 142,04 |
| HC_B1 | 0.0079 | 0.0073 | 24.45 | 93,27 |
| HC_B2 | 0.0066 |  |  | 97,78 |
| HC_B3 | 0.0079 |  |  | 104,34 |
| SC_B1 | 0.010 | 0.0147 | 26.78 | 124,40 |
| SC_B2 | 0.018 |  |  | 134,17 |
| SC_B3 | 0.015 |  |  | 107,51 |

### 3.4. Water absorption rate

Figure 11 shows the water absorption in relation to the time the specimen has been submerged. During the first 30 min. the samples had an initial similar and high water absorption rate, followed by a slower absorption. Only after 30 to 40 min. a linear trend appeared for SC and HC, which respectively lasted for 7 to 10 hours, after which the water absorption rate reduced. Consequently, the samples containing SC and HC absorbed water in a shorter period and reached their limit of absorption faster than FC. During the first 5 hours of the test the water absorption rate of FC and HC are similar. After 5 hours FC starts to absorb more water than HC. For the FC samples a non-linear behaviour appeared during the 24 hours of the test and the water absorption rate increased after 10 hours. Previously reported values by Ziegler et al., (2016) also displayed a non-linear nature of the water absorption curve [11]. This behaviour was assigned to the hydrophobic nature of mycelium and the hydrophilic nature of the fibres. The density of the samples can influence the diffusion coefficient of water. The low density of HC might affect the internal water transport. Moreover, as previously discussed the growth on HC samples resulted in a denser outer hydrophobic chitinous layer, compared to FC and SC, explaining the lower water absorption rate of HC.

**Figure 11:**
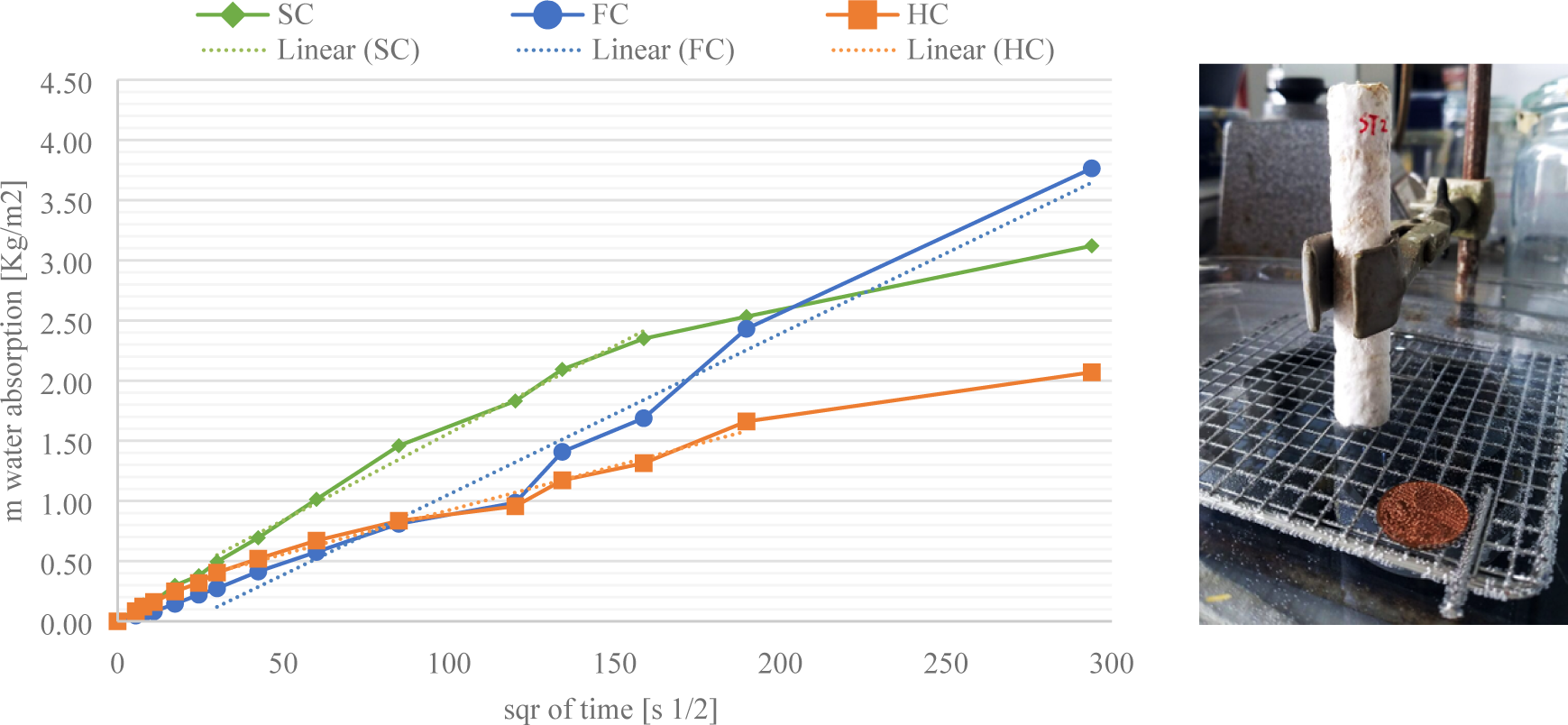
Plot of the water absorption of chopped flax (FC), hemp (HC) and straw (SC).

The velocity of HC (0,0073 mm/s^1/2^) samples to absorb water is lower than FC (0,0113 mm/s^1/2^) and SC (0,0147 mm/s^1/2^). After 24 hours FC had absorbed the highest amount of water (30,28%) compared to HC (24,45%) and SC (26,78%). A similar behaviour was observed with HC, the materials are not yet in a steady state after 24 hours and have the lowest absorption coefficient. Mycelium-composites made with HC are considered as interesting for construction purposes because of their low water absorption coefficient, and low absorption rate in 24 hours. All mycelium-composites have lower coefficients than clay bricks (0,019 mm/s^1/2^), mortar (0,011 mm/s^1/2^), glass fibres (0,049 mm/s^1/2^). The conducted tests could be improved in the future with a sensitivity analysis of the environmental conditions, different fibre sizes, homogeneity of the growth, and varying growth periods.

### 3.5. Chemical characterization

Since white-rot decay influences the chemistry of the fibres, a FTIR analysis was performed. *T. versicolor* is known to be a non-selective white-rot, it removes lignin and structural carbohydrates (hemicellulose, cellulose) at a similar rate [30]. White-rot fungi basidiomycetes are the most rapid degraders of lignin. By using phenol-oxidizing and peroxidase producing enzymes the fungi is able to depolymerise and mineralize lignin macromolecules [31]. The chemical structure of lignin therefore changes during the enzymatic oxidation with laccase and peroxidase [32,33]. During this oxidation process, lignin-based radicals can be cross-linked and subsequently form an adhesive between the fibres [13]. Spectra of all decayed and undecayed fibres presented in Table 1 were recorded, but in this paper we will only compare flax and hemp fibres, as previous discussed results for those fibres proved the most interesting advantages for construction applications. The peak position and intensities of the bands with respect to the baseline are listed in Table 6.

**Table 6:** Peak assignment for the FTIR characterization of *T. versicolor* grown on flax.

| Wave number (cm<sup>-1</sup>) | Peak (cm<sup>-1</sup>) | Assignment | Reference |
| --- | --- | --- | --- |
| 3600-3000 | 3280 | O-H stretching hydrogen bonds | [10,35] |
| 3000-2840 |  | C-H stretching in methyl and methylene groups | [30,36] |
| 2980-2835 | 2922, 2850 | CH<sub>2</sub>, CH<sub>2</sub>OH in cellulose |  |
| 2985-2947 | 2960 | CH<sub>2</sub> asymmetric stretching | [36] |
| 2940-2840 | 2850 | CH<sub>2</sub> symmetric stretching |  |
| 1770-1710 | 1733 | C=O stretching in xylans (hemicellulose) | [30,35] |
|  | 1633 | Absorbed O-H associated with lignin or cellulose | [30,36] |
| 1560-1520 | 1551 | C=C stretching of aromatic ring (syringyl) in lignin | [30,36] |
| 1520-1500 | 1510 | C=C stretching of aromatic ring (guaiacyl) in lignin | [30,35,36] |
| 1470-1410 |  | C-H deformation in methyl groups | Popescu |
|  | 1456, 1419 | C-H deformation in lignin and carbohydrates, CH<sub>2</sub> bending | [10,30,35,36] |
| 1375-1365 | 1370 | CH bending in cellulose and hemicellulose, chitin | [10,30,35,36] |
| 1365-1335 |  | OH plane deformation vibration | [36] |
|  | 1317 | CH<sub>2</sub> wagging in cellulose | [30,35,36] |
| 1282-1277 |  | CH deformation in cellulose | [36] |
|  | 1262 | Guaiacyl ring breathing, C-O linkage in guaiacyl aromatic methoxyl groups | [30,36] |
|  | 1240 | Syringyl ring, C-O stretching in lignin and xylan, Nuclei acids | [10,30,35] |
| 1235-1225 |  | OH plane deformation | [36] |
| 1205-1200 |  | OH plane deformation in cellulose, C-C stretching, C-O stretching, C-H deformation, | [10,36] |
| 1162-1125 | 1143 | C-O-C vibration in cellulose and hemicellulose | [30,35,36] |
|  |  | Aromatic C-H in-plane deformation | [36] |
|  | 1105 | Aromatic C-H in-plane deformation, C=O stretch | [36] |
| 1060-1015 |  | C-O valence vibration from C3-O3H | [36] |
| 1047-1004 |  | C-O stretching in cellulose | [30,35,36] |
|  | 1031 | C-C stretching | [10] |
|  | 896 | Anomeric C-group, Glucan β-anomer C-H bending, C-H deformation in cellulose | [10,30,35,36] |

The FTIR spectra of dried and undried flax composites are compared to undecayed fibres in Figure 12 (a), as well as the subtraction of undecayed peaks, and pure mycelium (without fibres) grown on flax. The subtracted peaks (long dash dot line) below the baseline show an increase or appearance of new bands due to the degradation by *T. versicolor*, while the subtracted peaks above the baseline show the decrease in intensity. The appearance and disappearance of peak after subtraction of decayed from undecayed fibres, clearly indicates new bonds created by the mycelium. The intensities are around zero when there is no change in the chemical composition of the fibres during the mycelium interaction.

**Figure 12:**
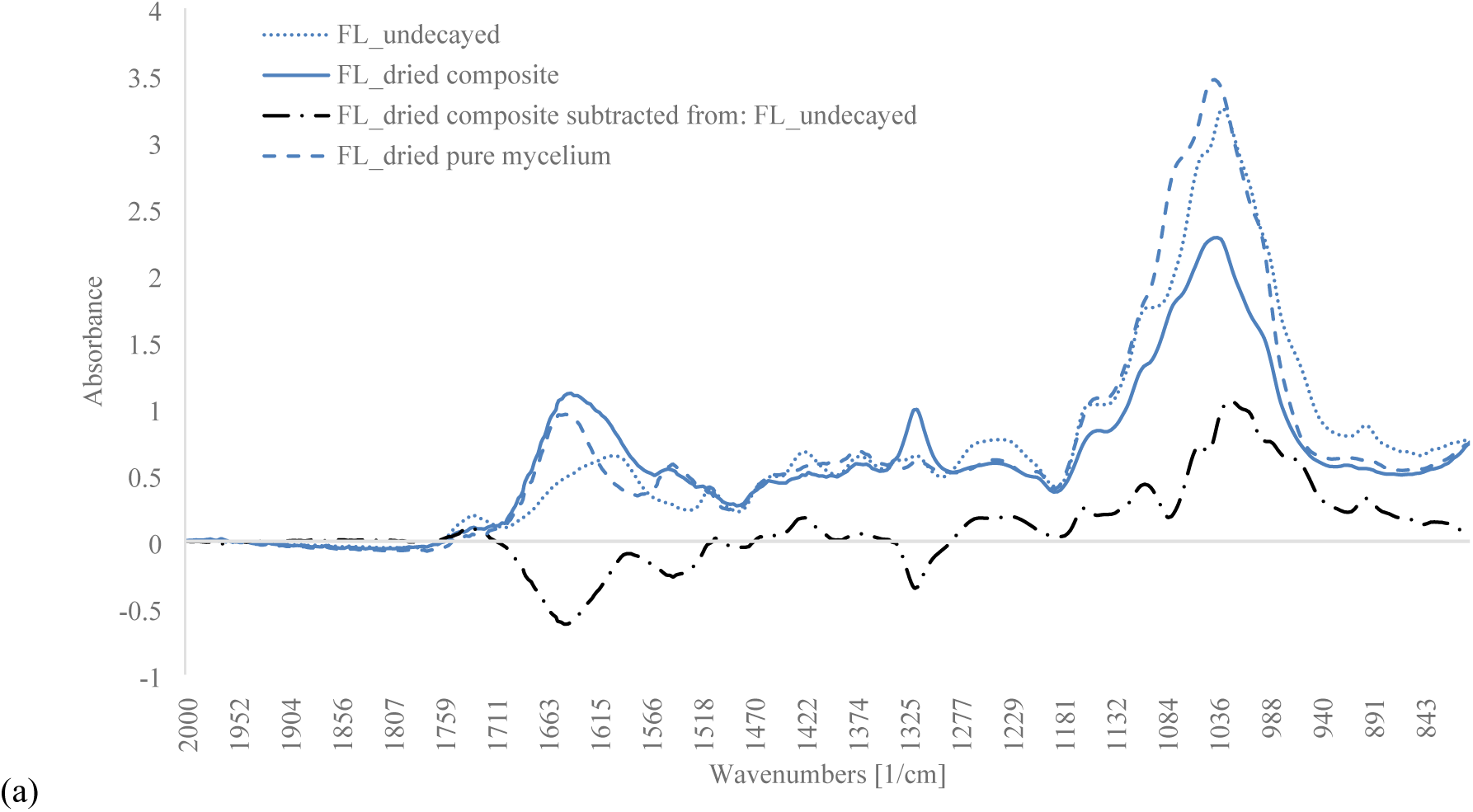
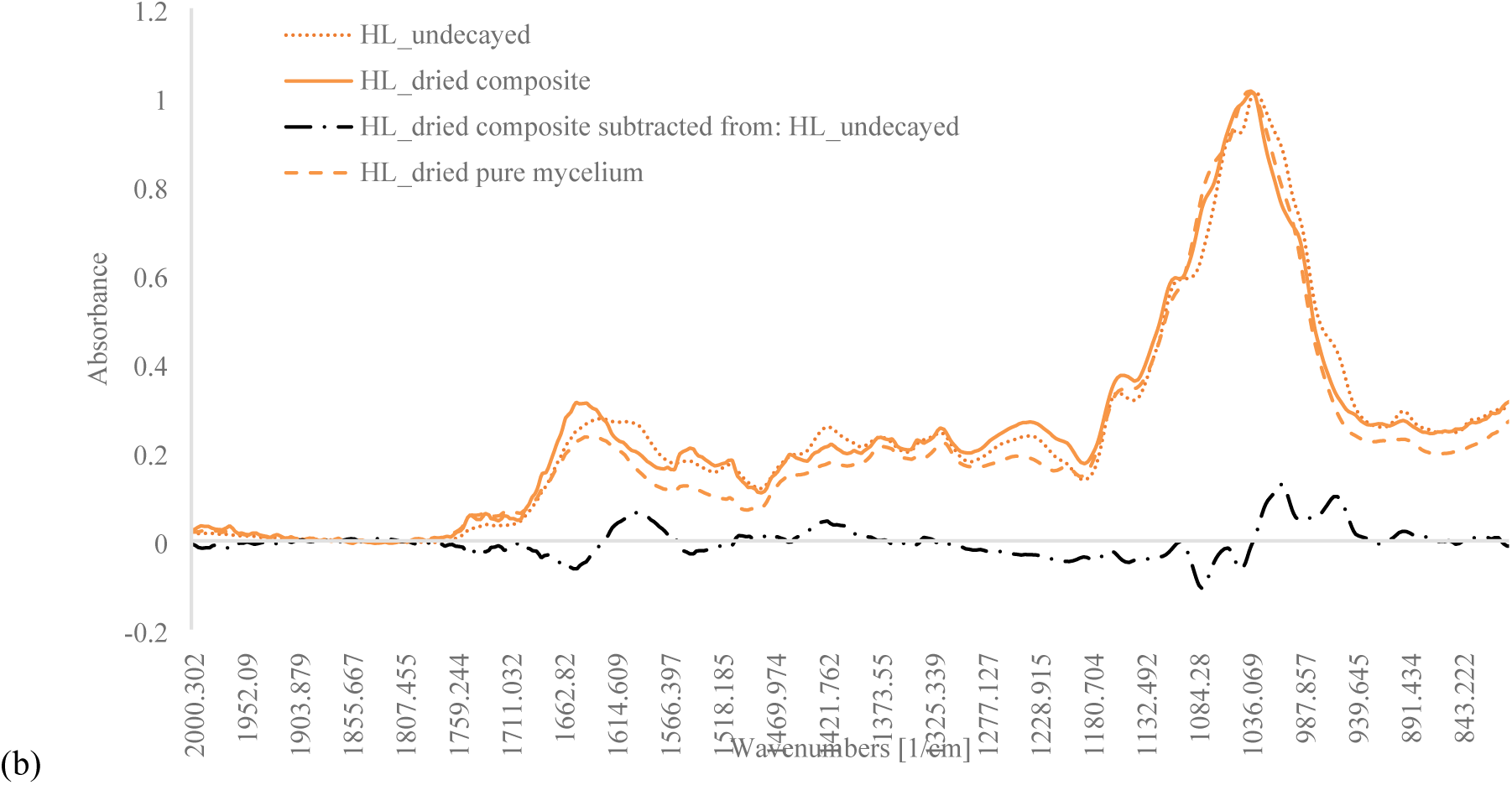
(a) FTIR spectra of undecayed loose flax fibres (solid line), undried loose flax composite (dotted line), dried loose flax composite (dashed line), and the subtraction dried composite from undecayed fibres (dot-dashed line). (b) FTIR spectra of undecayed loose hemp fibres (solid line), undried loose hemp composite (dotted line), dried loose hemp composite (dashed line), and the subtraction dried composite from undecayed fibres (dot-dashed line).

The degradation of the undried and dried flax composites by *T. versicolor* led to the small decrease in intensities of carbohydrates at 1733 cm^−1^ (weak), 1158 cm^−1^ (weak), 897.2 cm^−1^ (medium). The increase in carbohydrates was most pronounced at 1317 cm^−1^ (strong). The spectra also revealed an increase in the relative intensities of lignin bands at 1551 cm^−1^ (medium), 1510 cm^−1^ (strong), 1456 cm^−1^ (medium), and 1262 cm^−1^ (weak). Similar results are found for hemp, with a decrease carbohydrates intensities at 1733 cm^−1^ (weak), 1143 cm^−1^ (weak), 1047 cm^−1^ (weak), 896.8 cm^−1^ (medium). The degradation of hemp also resulted in an increase of lignin intensities at 1544 cm^−1^ (strong) and 1231 cm^−1^ (weak), accompanied by a decrease at 1595 cm^−1^, 1508 cm^−1^ (medium) and 1456 cm^−1^ (weak).

Figure 13 shows the comparison between the type of fibres by ratios of the relative intensities of lignin (1510 cm^−1^) against the carbohydrates (986 cm^−1^ and 1143 cm^−1^) for pure mycelium and dried composites. The ratio of lignin:carbohydrate intensities (I_1510_/I_986_ and I_1510_/I_1143_) between hemp and flax dried composites do not differ much. At first sight, this suggests that *T. versicolor* decays lignin, hemicellulose and cellulose in both fibre in the same way. The difference in ratios is more clear for pure mycelium hemp and flax samples. The ratios of peak intensities in lignin over cellulose and hemicellulose (I_1510_/I_1143_) exhibit a lower value (0,29) for hemp pure dried mycelium compared to (0,37) flax pure dried mycelium, which indicates a lower decay of lignin than cellulose in hemp compared to flax.

**Figure 13:**
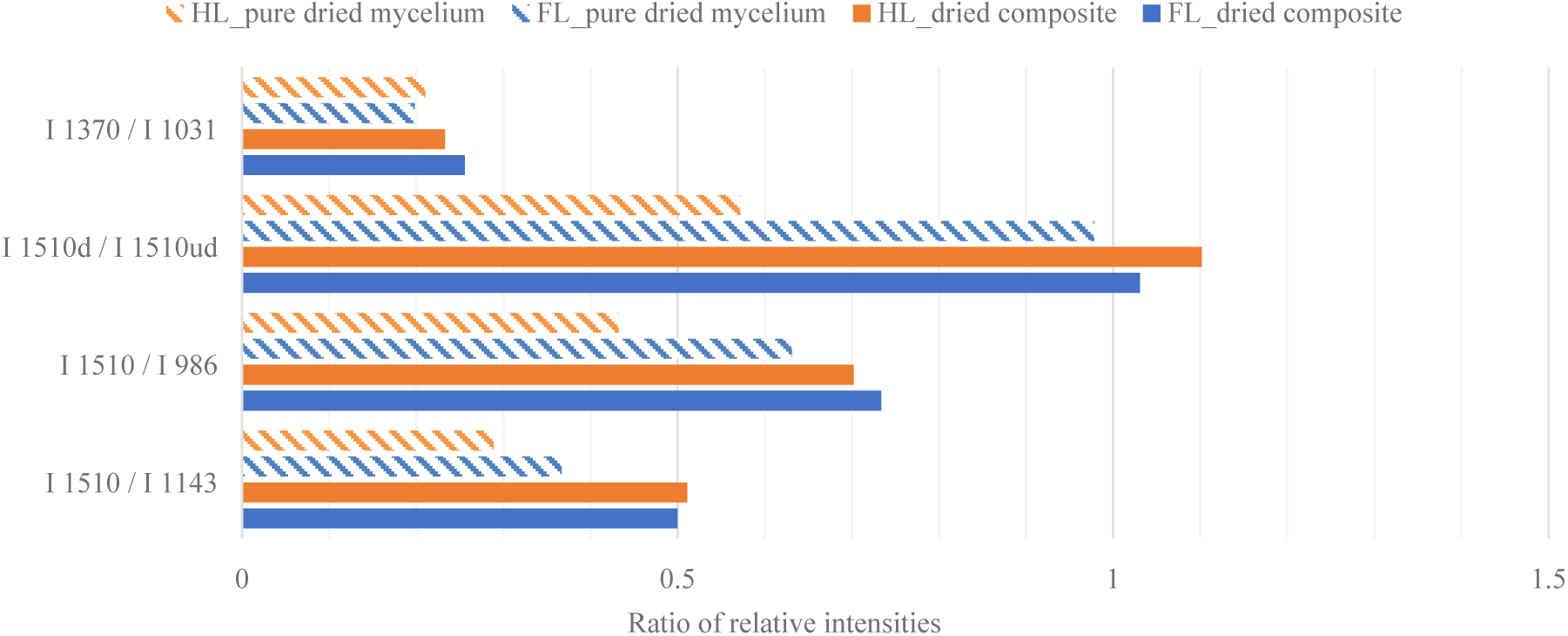
Ratios of the relative intensities of aromatic skeletal vibration in lignin I_1510_ against undecayed I_1510ud_ fibres, carbohydrate bands I_986_ (assigned to CH-deformation in cellulose) and I_1143_ (assigned to C-O-C vibration in cellulose and hemicellulose), C-H bending mode of chitin I_1370_ and the C-C stretching of polysaccharides I_1031_ of pure mycelium without fibres (diagonal stripes pattern), and dried hemp and flax composite (solid).

The decrease in the lignin:carbohydrate ratio, for hemp and flax, is higher for 1143 cm^−1^ band compared to 986 cm^−1^ band, indicating that *T. versicolor* has a small preference for hemicellulose over cellulose. Those results are consistent with other studies [30].

The relative presence of chitin was not influenced by the type of substrate, contrary to finding of Haneef et al. (2017). The ratio of peak intensity of chitin:polysaccharide (1370 cm^−1^ to 1031 cm^−1^) for hemp and flax was respectively 0,21 and 0,20 in pure mycelium, and 0,23 and 0,26 in the dried composites.

The ratio of peak intensities between decayed and undecayed fibres at 1510 cm^−1^ (aromatic skeletal vibration in lignin) are very similar for the composite fibres. The trend is more clear for pure mycelium, with a ratio of 0,57 for hemp and 0,97 for flax. The increase for flax reveals a higher intensity of lignin band of decayed flax than undecayed flax. The depolymerisation of lignin by *T. versicolor* happen to a greater extend in flax than in hemp. Previous studies have suggested that the decomposition of lignin, attributed to the phenol-oxidizing enzymes (laccase and peroxidase) affects adhesion and the composite’s strength. The _20_ enzymes promote the cross-linking of lignin-based radicals and increases the stiffness [34,37]. Furthermore, a high amount of cellulose in fibres results in a higher Young’s modulus and tensile strength [37]. Chopped flax fibres indeed showed a higher compressive stiffness (1,5 MPa) compared to chopped hemp (0,5 MPa), confirming previous research, but no consistency is found with the stiffness of loose fibres. Generally, the conducted tests reveal that the mechanical performances of the mycelium-based composites depend more on the fibre condition, size, processing, than on the chemical composition.

## 4. Conclusions

In this study we have investigated the requirements to produce mycelium-based composites with different types of lignocellulosic reinforcement fibres combined with a white rot fungi, *Trametes versicolor*. Together, they form an interwoven three-dimensional filamentous network binding the feedstock into a lightweight material. The mycelium-based material is heat-killed after the growing process. The purpose was to extend our knowledge of the mechanical, hygrothermal and chemical properties of mycelium-based composites. This is the first study reporting the dry density, the Young’s modulus, the compressive stiffness, the stress-strain curves, the thermal conductivity, the water absorption rate and a complete FTIR analyse of mycelium-based composites by making use of a disclosed protocol with *T. versicolor* and five different type of fibres (hemp, flax, flax waste, soft wood, straw) and fibre conditions (loose, chopped, dust, pre-compressed and tow).

This research showed that the production of mycelium-composites and their mechanical properties are dependent from the fibre types. Poor growth was found for dust flax and dust straw during the first and second growing period. This inconsistency may be due to unavailable nutrients and the absence of air-voids within the composite. Therefore, the fibre condition ‘dust’ was not taken into account for further testing. Wood- and straw-based composites also resulted in varying growth of *Trametes Versicolor*, consequently not all tests could be conducted with those fibres. A possible explanation for these results may be the difference in mould type for every test. The samples containing flax, hemp and flax-waste resulted in a well-developed composite and reliable results. Nevertheless, internal growth could be increased in further research by optimizing the fabrication procedure with sufficient air circulation in the mould.

Figure 14 shows the compressive stiffness of the various investigated samples. Overall, the results suggest that the fibre type (hemp, flax, wood, flax waste) has a smaller influence on the compressive stiffness than the fibre condition (loose, chopped, tow, pre-compressed). The Young’s modulus increased for all fibre types in chopped condition since the samples were more dense. Samples containing chopped fibres also resulted in a more coherent and smoother outer layer. As a compressive material, chopped flax-based samples obviously take the lead compared to the chopped hemp-based samples, which might have been affected by the manufacturing process. Furthermore, the pre-compression of the samples aimed to improve the compressive mechanical properties of mycelium composites. This fabrication method influenced the composite’s mechanical response in a positive way by increasing the Young’s moduli for every tested fibre type and condition (except for flax). These findings provide additional support for the hypothesis that compressed mycelium-composites enhance the mechanical performance, as presented in [9].

**Figure 14:**
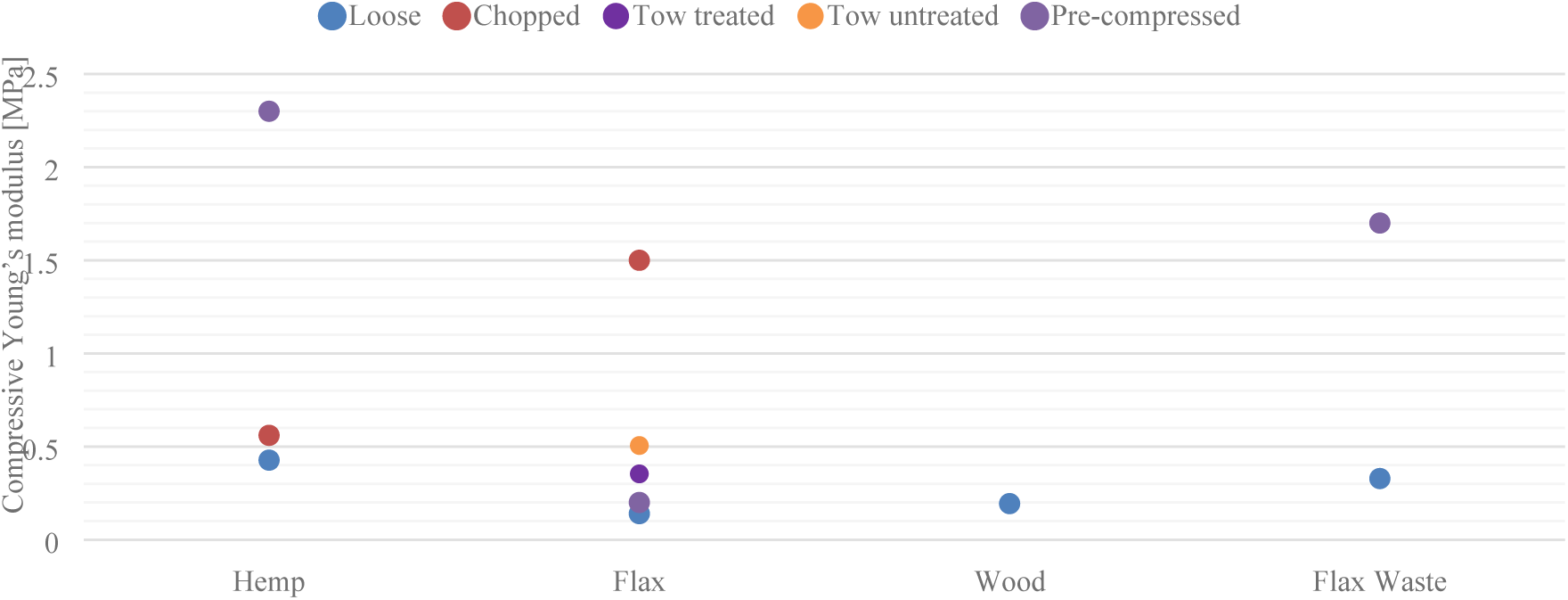
Comparision of the compressive strenght of different types of fibres (hemp, flax, wood and flax waste) in different fibre conditions (loose, chopped, tow treated, tow undtreated, pre-compressed).

Although the mechanical properties are not optimal yet, this research has shown that mycelium-composites can fulfil the requirements of thermal insulation. The thermal conductivity and water absorption coefficient of the mycelium composites with flax, hemp, and straw have shown an overall good insulation behaviour in all the aspects compared to conventional unsustainable materials (Figure 15). The hemp-based composites have the most interesting properties as an insulation material, with the lowest thermal conductivity 0.0404 [W/ (m*K)] and water absorption coefficient [0.0073 mm/s ^½^].

**Figure 15:**
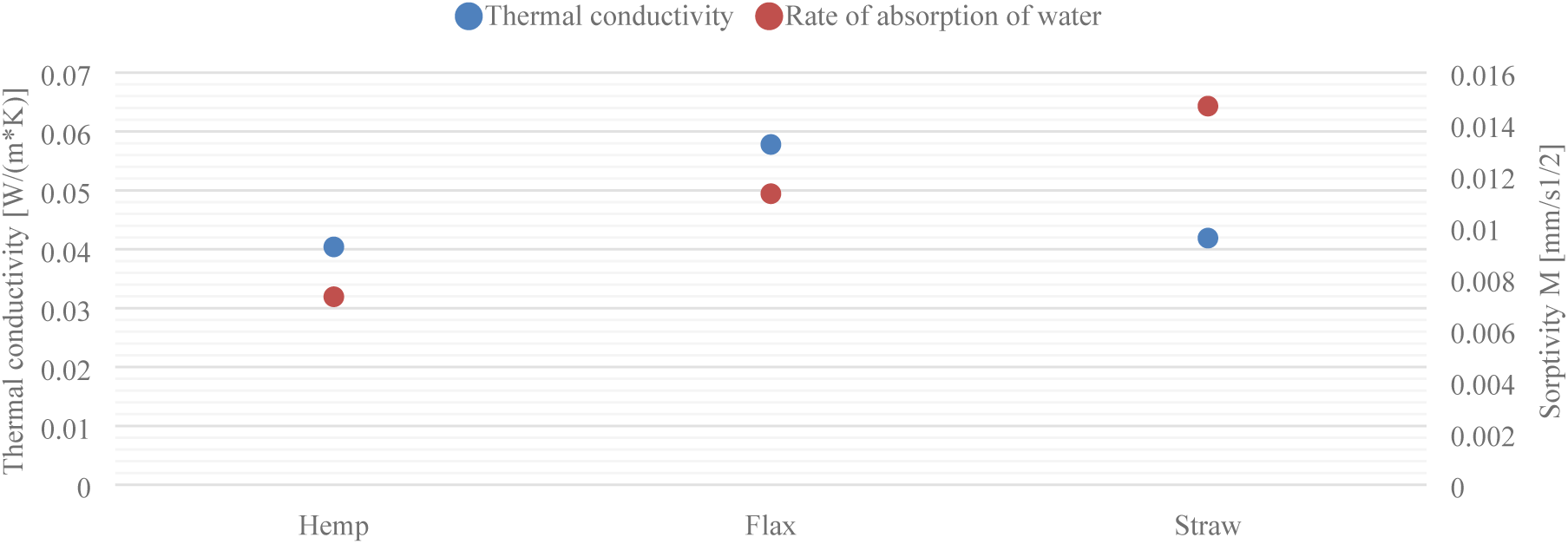
Comparison of the thermal and water absorption properties of different types of fibres (hemp, flax an straw).

The ratios of peak intensities of the FTIR spectra reveal a lower value for hemp pure dried mycelium in lignin over cellulose and hemicellulose compared to flax pure dried mycelium, which indicates a lower decay of lignin than cellulose in hemp compared to flax. The decrease in the lignin:carbohydrate ratio in hemp and flax indicated that *T. versicolor* has a small preference for hemicellulose over cellulose. The relative presence of chitin was not influenced by the type of substrate. The increase in ratio of peak intensities between decayed and undecayed flax fibres reveals a higher intensity of lignin band of decayed flax than undecayed flax. The depolymerisation of lignin by *T. versicolor* happen to a greater extend in flax than in hemp. Generally, the conducted tests reveal that the mechanical performances of the mycelium-based composites depend more on the fibre condition, size, processing, than on the chemical composition.

The wide spectrum of options to compose and grow mycelium-composites makes it complex to compare the results with existing literature since every change of variable will affect the growth and mechanical behaviour of the composite. Further work is required to improve the growth conditions, to optimize the mechanical properties and to establish a standard fabrication protocol.

## Acknowledgements

The authors would like to thank the master-thesis students Aurélie Van Wylick and Paola Viviana Pantoja Arboleda for their dedicated contributions during this research.

## Data availability

All processed and raw data files are available from the Mendeley Data database (accession number doi:10.17632/r8m4y7rv8j.1)

